# Strain-dependent modifier genes determine survival in *Zfp423* mice

**DOI:** 10.1101/2020.05.12.091629

**Authors:** Wendy A. Alcaraz, Zheng Liu, Phoebe Valdes, Edward Chen, Alan G. Valdovino Gonzalez, Shelby Wade, Cinny Wong, Eunnie Kim, Hsiang-Hua M. Chen, Alison Ponn, Dorothy Concepcion, Bruce A. Hamilton

## Abstract

*Zfp423* encodes a transcriptional regulatory protein that interacts with canonical signaling and lineage pathways. Mutations in mouse *Zfp423* or its human ortholog *ZNF423* are associated with a range of developmental abnormalities reminiscent of ciliopathies, including cerebellar vermis hypoplasia and other midline brain defects. Null mice have reduced viability in most strain backgrounds. Here we show complete lethality on a C57BL/6J background, dominant rescue in backcrosses to any of 13 partner strains, with strain-dependent survival frequencies, and evidence for a BALB/c-derived survival modifier locus on chromosome 5. Survival data indicate both perinatal and postnatal periods of lethality. Anatomical data from a hypomorphic gene trap allele observed on both C57BL/6J and BALB/c congenic backgrounds shows an aggregate effect of background on sensitivity to Zfp423 loss rather than a binary effect on viability.

## INTRODUCTION

Zfp423 and its human homolog ZNF423 are transcriptional regulators that intersect both lineage and signal-induced pathways. Zfp423 was first described as an inhibitory interacting factor for Ebf-family basic helix-loop-helix (bHLH) transcription factors, blocking Ebf-dependent development in olfactory neurogenesis (Cheng and Reed 2007; Tsai and Reed 1997, 1998). Zfp423 was later independently found as a binding partner of BMP-dependent SMAD transcription factors (Hata et al. 2000), retinoic acid receptors (Huang et al. 2009), cleaved Notch intracellular domain (Masserdotti et al. 2010) and DNA damage response proteins (Chaki et al. 2012). Recruitment of Zfp423 in one pathway appears to antagonize alternative pathways (Hata et al. 2000; Masserdotti et al. 2010), suggesting that Zfp423 may act to integrate competing signals into a canalized cellular response. Indeed, Zfp423 is a potent factor for embryonic stem cell exit from naïve state (Lackner et al. 2020) and Zfp423 architecture and sequence are highly conserved (Hamilton 2020).

*Zfp423*-mutant mice have characteristic developmental abnormalities, particularly in hindbrain. Loss of cerebellar vermis and other midline defects in developing brain are prominent features across independent null and hypomorphic alleles (Alcaraz et al. 2006; Cheng et al. 2007; Warming et al. 2006). Vermis hypoplasia originates in reduced numbers of proliferating progenitors adjacent to the midline (Alcaraz et al. 2006) and Zfp423 in cerebellar granule cell progenitors (GCPs) is required for mitogenic response to SHH signaling, potentially through repression of Tulp3 (Hong and Hamilton 2016). Development of the roof plate and choroid plexus in the fourth ventricle are frequently abnormal (Alcaraz et al. 2006; Casoni et al. 2020). Formation of hippocampus, septum and thalamus are often affected, as are several fiber tracts, including frequent loss of midline crossing by commissural axons in corpus callosum and reduction of the anterior commissure (Alcaraz et al. 2006; Cheng et al. 2007). Loss of Zfp423 impairs maturation of olfactory neurons in a manner consistent with it acting in a developmental switch between early differentiating and maturing olfactory neurons (Cheng and Reed 2007). Zfp423 is also expressed by preadipocyte fibroblasts, where it induces PPARg-dependent adipogenesis, and loss of *Zfp423* function impairs adipogenesis both in development (Gupta et al. 2010) and in wound healing (Plikus et al. 2017).

Mutations in the human homolog *ZNF423* have been reported in patients with ciliopathy diagnoses and anatomical findings reminiscent of *Zfp423* mutant mice (Chaki et al. 2012) and a subset of patient mutations has been validated in mouse models (Deshpande et al. 2020). Ciliopathies comprise a spectrum of related phenotypes that involve one or more organs, often cerebellum, kidney, retina, and/or liver, and are united by cellular defects in structure of and signaling through the primary cilium (Badano et al. 2006; Hildebrandt et al. 2011; Waters and Beales 2011). Ciliopathies also range in severity from nearly unaffected to prenatal lethal. Some of the differences arise from differences between causal genes or allele strength, but similar primary mutations can give rise to very different outcomes in both patients and mouse models depending on other genes and non-genetic factors (Davis and Katsanis 2012; Norris and Grimes 2012). In previous work, we showed that differences in cerebellar hypoplasia had both genetic and non-genetic components in BALB/c and 129S1 strains (Alcaraz et al. 2011), but the distribution and effect sizes of genetic modifiers and other sources of variability are not well explored.

Here, we show complete lethality of *Zfp423* mutants in a C57BL/6 (B6 hereafter) background that correlates with both adiposity and neonatal feeding, and that lethality is specific to the perinatal period. As a prologue to both modeling patient mutations in mice and identifying modifiers that influence Zfp423-deficient developmental outcomes, we surveyed 13 inbred mouse strains through a series of (B6–*Zfp423*^*nur12*^ x strain) x B6–*Zfp423*^*nur12*^ backcrosses. Surprisingly, even C58, C57L, and C57BLKS, strains closely related to B6, were able to support a modest number of surviving pups, some to a year or more of age. Calculated survival rates range from 8% to 95% depending on the strain pairing. Legacy data from earlier B6–*Zfp423*^*nur12*^ x BALB/c intercross and backcross mice allowed us to infer a modifier locus on chromosome 5, explaining a fraction of survival differences between those strains.

## MATERIALS AND METHODS

### Mice

BALB/cAnNHsd (BALB/c) mice were obtained through the UC San Diego Transgenic Mouse Shared Resource and maintained in our laboratory. PWK/PhJ mice were obtained from Dr. Christopher K. Glass. 129S1/SvImJ (129S1), A/J, BTBR T/+ tf/tf (BTBR), C3H/HeJ, C57BL/6J (B6), C57BLKS/J, C57L/J, C58/J, DBA/2J (DBA), FVB/NJ (FVB), NOD/LtJ, and NZO/HlLtJ were obtained from The Jackson Laboratory. Congenic *Zfp423*^*nur12*^ and *Zfp423*^*XH542*^ mice were developed by serial backcross from mixed stock and linked markers on chromosome 8 were followed in early generations to minimize linked variation as previously described (Alcaraz et al. 2011; Alcaraz et al. 2006) and maintained by further backcross in our laboratory. Crosses described here began with fully congenic stocks for B6–*Zfp423*^*nur12*^ (N28, with no detected heterozygosity among 24 polymorphic markers on chromosome 8), BALB/c–*Zfp423*^*nur12*^ (N31, with a congenic interval ~2.4 Mb bounded by *D8Mit57* and *D8Mit327*), and 129S1–*Zfp423*^*nur12*^ (N24 without further characterization of the interval after Alcaraz et al. 2011). Mice were maintained on LabDiet 5P06 chow ad libitum and a 12 hr light:12 hour dark cycle in microisolator cages on high-density racks in a specific pathogen free facility, but crosses were split between two facilities on the UC San Diego campus, spanning a lab relocation. All animal experiments were approved by the UC San Diego Institutional Animal Care and Use Committee (IACUC).

### Crosses

To test survival on different fully inbred or hybrid backgrounds, congenic stock heterozygotes were bred inter se to produce congenic F1 or crosses between congenic lines to produce hybrid F1 offspring. For 129S1, A/J, BALB/c, and FVB, crosses were initiated in each direction, with B6 as either the female or male partner, with equivalent results. For ease of breeding, other strain pairs were initiated with two females of the inbred strain crossed to males of the B6-*Zfp423*^*nur12*^ congenic stock. Backcrosses to B6-*Zfp423*^*nur12*^ congenic stock were performed using F1 of either sex, depending on availability of B6-*Zfp423*^*nur12*^ congenic and F1 breeders. Unless otherwise noted, survival was scored at two weeks postnatal. Where sufficient numbers were available for retrospective tests, no differences in survival frequency were detected between backcross of F1 hybrid females and F1 hybrid males of a given strain pair and data from each direction were analyzed as a single cross. Statistical comparisons were made in R 3.5.1.

### Histology

P0 pups were fixed in 10% buffered formalin, cryoprotected with 30% sucrose, and frozen in OCT before being sectioned at 20 μm. Mice with the Zfp423 gene trap allele XH542 were sacrificed under deep anesthesia and fixed by transcardial perfusion with saline and 4% paraformaldehyde at 2 weeks old, cryoprotected with 30% sucrose, and frozen in OCT and sectioned at 16 μm. Cryosections were stained with hematoxalin and eosin. Sections were visualized and photographed on a Nikon E800 microscope equipped with a Spot Xplorer digital camera under standard illumination. Measurements were made from .tif format files in ImageJ.

### Linkage analysis

Simple sequence-length repeat marker typing was performed using standard methods (Alcaraz et al. 2006; Dietrich et al. 1996). Marker loci were tested for reduced frequency of B6 allele homozygosity among surviving *Zfp423*^*nur12*^ homozygous pups using a chi-squared test in the R base environment from genotype frequencies tabulated in R/qtl (Broman and Sen 2009; Broman et al. 2003). Tests for duration of survival, up to 60 days, was carried out using the 2-step procedure in the scanone function and proportion of variance explained was estimated using the fitqtl function in R/qtl (Boyartchuk et al. 2001; Broman 2003).

### Data Availability Statement

All data are included in the figures and tables or in supplemental tables for genotyping data. *Zfp423*^*nur12*^ mice are available from the Mouse Mutant Regional Resource Centers.

## RESULTS

### *Zfp423* null allele is perinatal lethal in B6 strain background

We examined *Zfp423*^*nur12/+*^ mice as congenic lines on three common inbred strain backgrounds and as hybrids between them. Offspring were genotyped at 1 to 2 weeks. While *Zfp423*^*nur12/nur12*^ showed somewhat reduced frequency in BALB/c and 129S1 congenic strains, *Zfp423*^*nur12/nur12*^ mice were never recovered among the B6 congenic strain (Table 1). When B6–*Zfp423*^*nur12/+*^ mice were crossed to BALB/c–*Zfp423*^*nur12/+*^ mice, the frequency of *Zfp423*^*nur12*^ homozygotes was consistent with full survival. The difference in homozygote frequency between BALB/c congenic stock and the BALB/c x B6 F1 (*P* = 0.019, Fisher’s exact test) suggests that both strains contribute recessive alleles that reduce *Zfp423*^*nur12/nur12*^ viability.

**Table 1:**
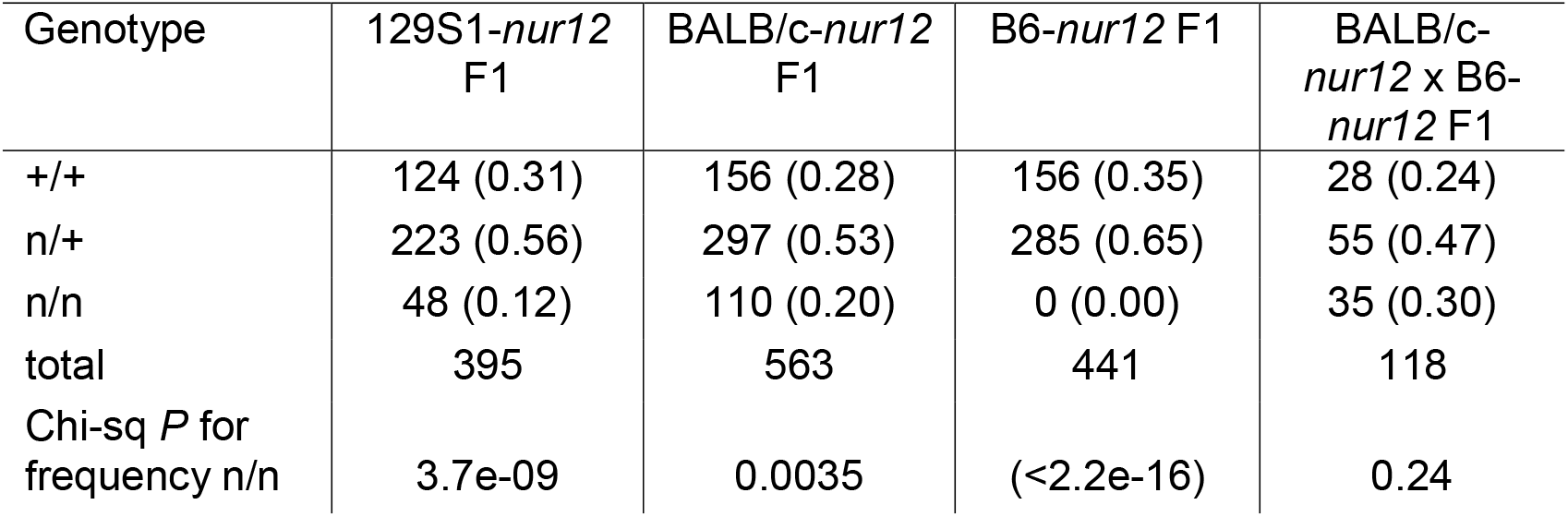
*Zfp423*^*nur12*^ genotype frequencies (proportions) show enhanced lethality in B6 background.

| Genotype | 129S1-*nur12*<br>F1 | BALB/c-*nur12*<br>F1 | B6-*nur12* F1 | BALB/c-*nur12* x B6-*nur12* F1 |
| --- | --- | --- | --- | --- |
| +/+ | 124 (0.31) | 156 (0.28) | 156 (0.35) | 28 (0.24) |
| n/+ | 223 (0.56) | 297 (0.53) | 285 (0.65) | 55 (0.47) |
| n/n | 48 (0.12) | 110 (0.20) | 0 (0.00) | 35 (0.30) |
| total | 395 | 563 | 441 | 118 |
| Chi-sq *P* for frequency n/n | 3.7e-09 | 0.0035 | (<2.2e-16) | 0.24 |

To determine lethal phase and rule out segregation distortion or gametic effects, timed embryos and postnatal pups were collected and genotyped to determine the frequency of *Zfp423*^*nur12*^ homozygotes at each age. Only two mutants were observed at P1 and neither survived to P2. These data show complete perinatal lethality of *Zfp423*^*nur12*^ homozygotes on the B6 congenic background (Table 2).

**Table 2:** Genotype frequencies (proportions) by age support perinatal lethality.

| genotype | E12.5 | E14.5 | E16.5 | E18.5 | P0 | P1 |
| --- | --- | --- | --- | --- | --- | --- |
| +/+ | 27<br>(0.20) | 25<br>(0.21) | 17<br>(0.22) | 35<br>(0.26) | 59<br>(0.29) | 22<br>(0.30) |
| n/+ | 76<br>(0.55) | 62<br>(0.53) | 40<br>(0.51) | 77<br>(0.57) | 108<br>(0.54) | 49<br>(0.67) |
| n/n | 34<br>(0.25) | 31<br>(0.26) | 21<br>(0.27) | 23<br>(0.17) | 33<br>(0.16) | 2<br>(0.027) |
| total | 137 | 118 | 78 | 135 | 200 | 73 |
| Chi-sq *P* for frequency n/n | 0.96 | 0.75 | 0.69 | 0.032 | 0.005 | 1.2e-05 |
| Fisher's exact *P* for comparison to previous day | - | 0.89 | 1 | 0.11 | 0.76 | 0.0017 |

### Reduced feeding and brown adipose tissue in B6–*Zfp423* neonates

To identify possible causes of lethality in B6-*Zfp423*^*nur12/nur12*^ mice, we examined histology of P0 pups (Figure 2). Heart, kidney, and liver did not show any obvious defects. Sections through stomach show reduced amount of milk in B6 compared to BALB/c mice, independent of *Zfp423* genotype, while *Zfp423*^*nur12/nur12*^ mice have reduced amount of milk compared to littermate controls in each strain, such that B6–*Zfp423*^*nur12/nur12*^ mice are least well fed among comparison groups. Examination of brown adipose tissue confirmed small fat pads with gaps in the B6-*Zfp423*^*nur12/nur12*^ mice as expected (Gupta et al. 2010). Two out of the three BALB/c-*Zfp423*^*nur12/nur12*^ mice examined have normal brown adipose appearance, however, suggesting a degree of strain-dependence for this phenotype in limited sampling.

**Figure 1.**
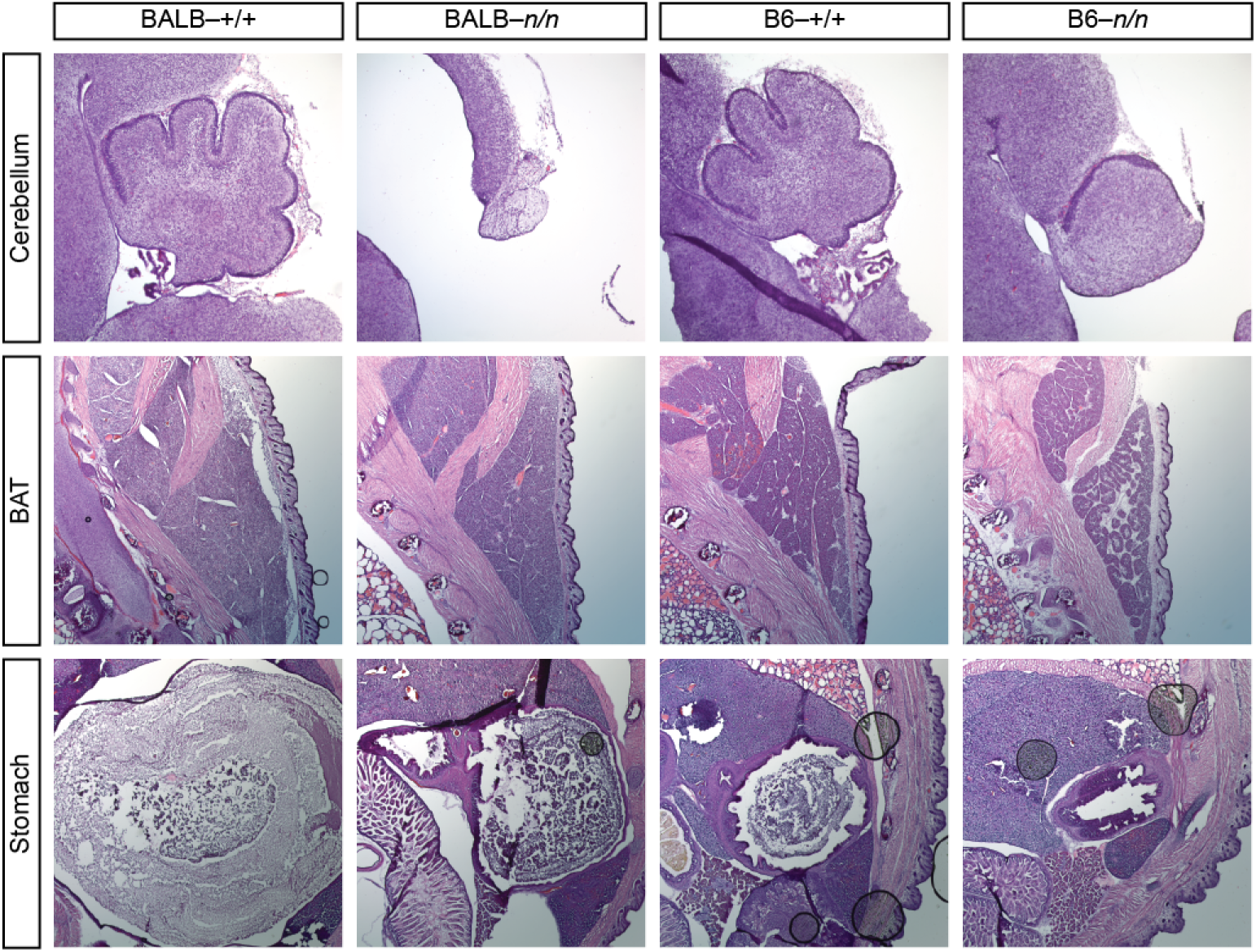
Histological differences between strains at P0. Typical images from a small series of P0 animals. *Zfp423*^*nur12/nur12*^ homozygotes on both BALB/c and B6 backgrounds showed cerebellar vermis hypoplasia, anterior rotation and reduction or absence of choroid plexus. Brown adipose tissue generally appeared intact on BALB/c but reduced and less compact on B6. Stomach content was reduced in mutants, especially on B6 background.

**Figure 2.**
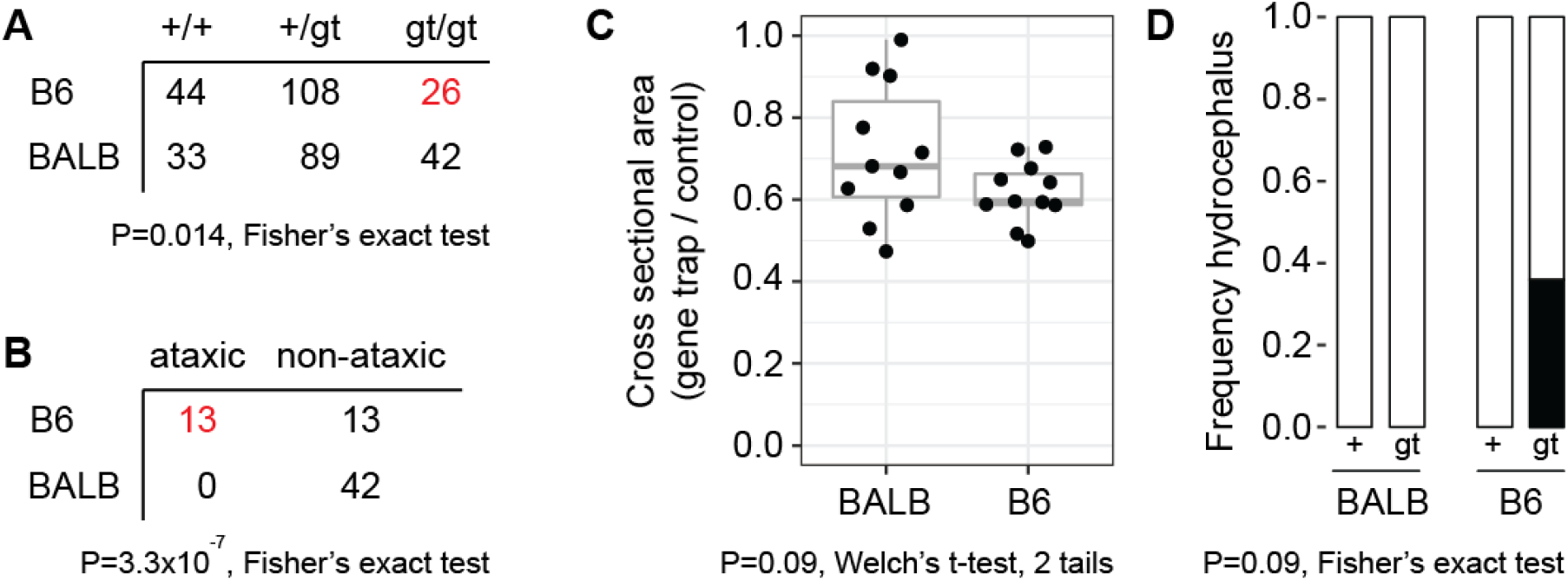
Congenic gene trap lines support B6 sensitivity to reduced Zfp423 function. (A) Gene trap (gt) homozygotes are recovered with reduced frequency on B6 but not BALB/c congenic background. (B) Surviving B6-gene trap homozygotes were frequently judged ataxic prior to genotyping, while BALB/c-gene trap homozygotes were not. (C) Parasagittal sections through cerebellar vermis showed reduced area in gene trap homozygotes compared to littermates, but the trend toward larger difference on B6 was not significant at alpha = 0.05 level for a two-tailed test. (D) A subset of B6 gene trap homozygotes were hydrocephalic, while none of the other mice were.

### B6 background enhances phenotype of a *Zfp423* hypomorphic allele

To determine whether strain background might similarly affect weaker *Zfp423* alleles, we examined congenic lines for a hypomorphic gene trap line, XH542, that expresses ≤50% normal *Zfp423* RNA levels (Alcaraz et al. 2006), on both BALB/c and B6 backgrounds (Figure 2). Gene trap homozygotes are viable on both backgrounds, but occur with reduced frequency on B6 (15% vs 26%, Figure 2A). Surviving B6 gene trap homozygotes also have higher frequency of ataxia (Figure 2B). Gene trap homozygotes on both strain backgrounds had reduced cross-sectional area in vermis, but the difference between strains was not significant at conventional alpha for pre-planned sample size (N=11 paired samples, Figure 2C). Among samples allocated to anatomy we observed four with hydrocephalus, all of which were B6 gene trap homozygotes (Figure 2D). Taken together, these observations further support B6 as a sensitized background for loss of *Zfp423* function.

### B6 sensitivity to *Zfp423* relative to BALB/c is polygenic, recessive in aggregate, and includes a locus on chromosome 5

To test the genetic architecture of the striking difference in *Zfp423* survival and phenotype severity, we analyzed data from both B6 x BALB/c intercross and backcross designs performed several years ago (Supplemental Data S1 and S2). We specified linkage thresholds for suggestive (expected once per genome by chance) and significant (once in twenty genomes by chance) for a given cross design in advance (Lander and Kruglyak 1995) and assessed both marker-based chi-squared tests for proportion of B6 homozygotes at ascertainment (~2 weeks postnatal) and for survival up to P60 as a quantitative trait. In the intercross, chi-squared tests for proportion of B6 homozygotes (1 degree of freedom) supported linkage at genome-wide significance for reduced B6 homozygotes at a locus on chromosome 5 (*P* = 3.1 × 10^−4^ at rs254857227), with significant over-representation of B6 homozygotes on chromosome 13 (*P* = 3.7 × 10^−4^ at *D13Mit17*; Figure 3A). Similar analysis of an earlier intercross from a partially inbred stock had shown additional strong signals on chromosomes 11 and 17 that were not replicated in the intercross from fully congenic stocks, although the chromosome 17 locus remained significant in a combined analysis. This suggested context dependence and potential for epistatic interactions and resulted in limiting formal analysis to crosses initiated from fully congenic stocks. Analysis of a smaller backcross showed only suggestive levels of support for a locus on chromosome 12 (*P* = 0.0012) and sub-threshold support for the locus on chromosome 5 (*P* = 0.01), but encompassing the same chromosome 5 interval as the intercross (Figure 3B). Selective genotyping of ~70 backcross progeny from other strain partners (see below) for survival at two weeks increased support for the chromosome 5 locus, with combined support across all backcrosses in the suggestive range (*P* = 6.9 × 10^−4^ at *D5Mit157*) for backcross data alone.

**Figure 3.**
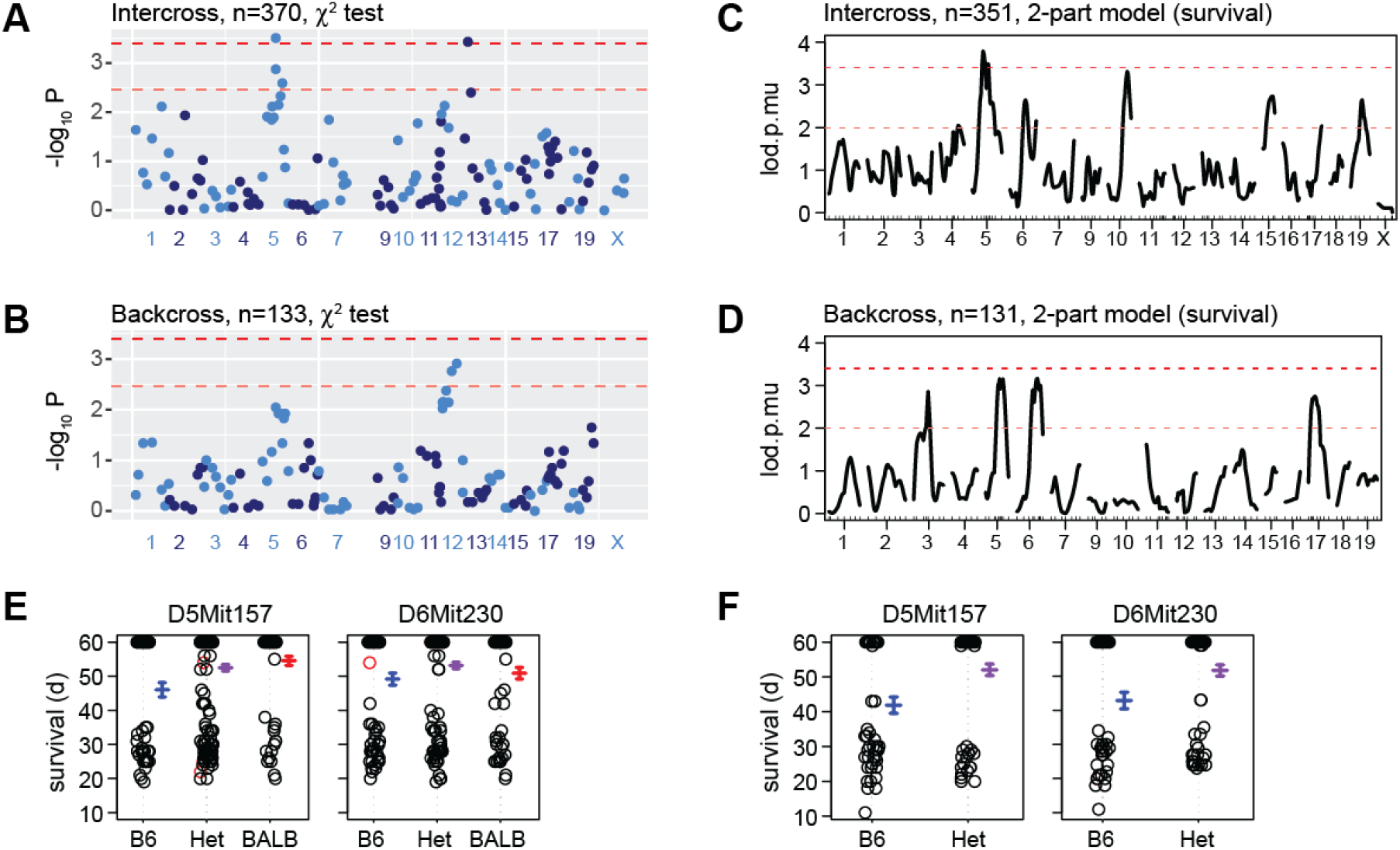
Linkage mapping supports a survival locus on Chr. 5. (A) Chi-squared tests for expected 25% B6-homozygous fraction (1 degree of freedom) in 370 F2 progeny from a BALB/c x B6 intercross. Chromosome 8 is omitted as constrained by *Zfp423* requirement. (B) Chi-squared test for 50% homozygotes in 133 backcross progeny. (C) 2-part model for survival (latency) up to P60 for the intercross progeny in (A). Lod.p.mu is the combined lod score for penetrance (lod.p) and severity (lod.mu) using log survival from entry at P12 to endpoint at P60 as a quantitative phenotype. (D) 2-part model applied to the backcross progeny in (B). (E) Marker effect plots for survival up to postnatal day (d) 60 in the intercross. (F) Effect plots for the same markers in the backcross.

Most animals from both intercross and backcross populations also had survival data to postnatal day 60, which allowed use of a 2-part model for survival duration as a quantitative trait (Boyartchuk et al. 2001; Broman 2003). Analysis of the intercross again showed significant linkage to the middle of chromosome 5, while also finding suggestive linkage on several other chromosomes (Figure 3C). Similar analysis of survival data from the backcross showed suggestive support for four regions, with strongest support for the chromosome 5 locus. In addition, the backcross animals that survived to at least P60 had significant depletion of B6 homozygotes compared to heterozygotes (22 vs. 59, chi-squared *P* = 3.9 × 10^−5^ at *D5Mit398*-*D5Mit157*) and was the most significant finding genome-wide for survival to P60. Two-part survival analysis combining the intercross and backcross data (for loci that might act independently in both contexts) showed strongest support was for the chromosome 5 locus (lod.p.mu = 7.4) with significant support for a locus on chromosome 6 (lod.p.mu = 5.1). Both loci were significant at genome-wide alpha < 0.02 based on 5000 permutations of the data. Effect plots for intercross (Figure 3E) and backcross (Figure 3F) showed that both loci reduce the probability of death at 3-5 weeks in favor of survival to the censored time point at P60. A simple additive model for the loci on chromosomes 5 and 6 applied to the combined backcross and intercross data explained 10% of the variance in survival, with a LOD score of 11.6 and most of the estimated effect contributed by chromosome 5 (6.5%).

### B6 synthetic lethality is recessive and rescued by any of 13 strains

To examine the viability effects of strain background more broadly and to identify strains that might be useful for testing patient variants in an animal model, we crossed B6–*Zfp423*^*nur12/+*^ to broad panel of inbred strains (Figure 4A) and then backcrossed the resulting F1 progeny to B6–*Zfp423*^*nur12/+*^ to produce F2 (Figure 4B). Crosses with each of the 13 partner strains produced viable mutant offspring, regardless of which sex was used as the F1 for backcross. The survival difference can not, therefore, be explained by B6 maternal behavior. Progeny of each cross were genotyped for *nur12* and all identified homozygotes had clearly recognized gait ataxia, indicating full penetrance. While some strains were under-sampled due to poor breeding performance, clear differences emerged among partner strains (Figure 4C). The frequency of F2 mutants recovered from the PWK/PhJ cross was 24% and not significantly different from Mendelian expectation (*P* = 0.84, chi-squared test, 1 df). By contrast, every other backcross showed reduced recovery, from 13% in 129S1 (*P* = 2.8 × 10^−5^), to 2.5% in BTBR (*P* = 1.5 × 10^−8^). The difference among strains had strong statistical support (*P*=5.0 × 10^−4^, Fisher’s exact test; support for pairwise comparisons are provided in Ssupplemental Table S1). These data showed that survival of *Zfp423*^*nur12*^ homozygous mice is polygenic, with different sets of survival factors in different inbred stains.

**Figure 4.**
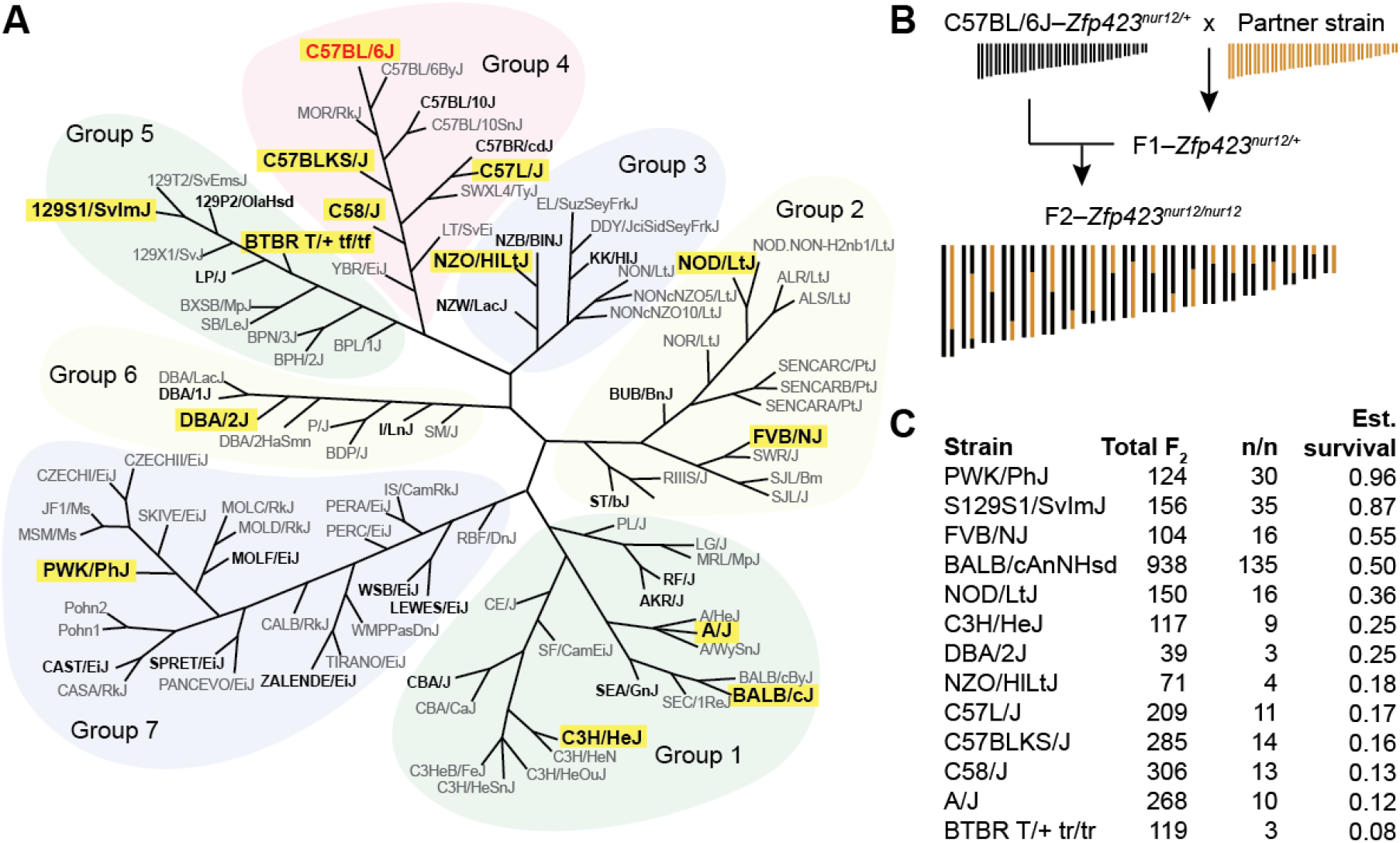
Thirteen diverse strains rescue B6– *Zfp423*^*nur12/+*^ lethality. (A) SNP-derived neighbor-joining tree modified from Petkov et al. (Petkov et al. 2004). Strains with public genome sequences (Keane et al. 2011) are in bold, strains tested for rescue of B6-dependent lethality highlighted with yellow rectangle. Only B6 (C57BL/6J, shown in red) showed complete lethality. (B) Outcross-backcross design tests for dominant (or semi-dominant) effects. (C) Survival fraction was estimated as the number of *Zfp423*^*nur12*^ homozygotes divided by one-third the number of non-mutant offspring from the same cross.

## DISCUSSION

Our results showed that the *Zfp423*^*nur12*^ null allele is a fully penetrant perinatal lethal on a C57BL/6J (B6) genetic background, but has ≥50% survival when congenic on either 129S1 or BALB/c. Of the few neonates available for examination, mutants on the B6 background appeared to have more severe peripheral tissues abnormalities compared with mutants congenic on BALB/c, including notably reduced adipose tissue and poor feeding. These observations are consistent with previous reports on Zfp423 requirements for adipocyte differentiation (Gupta et al. 2010; Gupta et al. 2012; Plikus et al. 2017; Shao et al. 2017) and suggest that this phenotype in vivo may be strain-dependent in its effect size. Greater sensitivity of B6 than BALB/c for loss of *Zfp423* function was supported by a small series of congenic mutants for a hypomorphic gene trap allele. Both survival and behavioral ataxia were substantially more penetrant on B6 than BALB/c, although statistical support for anatomical differences were sub-threshold and likely underpowered relative to the observed variance. This strain difference in sensitivity contrasts with recent work in our lab comparing different alleles on a single inbred background, where anatomical differences were more sensitive than behavior in a more extensive series of animals (Deshpande et al. 2020). Surviving *Zfp423*^*nur12*^ mutants from intercross and backcross designs reported here showed that no single locus is either necessary or sufficient for mutant survival, but taken together, our results supported the existence of a modifier locus on chromosome 5, with suggestive or significant support for linkage at other loci being cross-dependent. Comparing survival rates among B6 backcross progeny from several different partner strains indicated different sets of pro-survival factors in different inbred strains. This suggests that searches for survival loci might be more powerful in strains with low survival frequency and close relation to B6, as these should limit the number of contributing loci and therefore the statistical power per surviving mutant. These results also informed our choice of strains for modeling patient variants of uncertain significance (Deshpande et al. 2020), where FVB provided a high-survival, yet fully-penetrant background compatible with recovery of severe alleles and B6 provided a sensitized background to increase confidence in negative findings (Deshpande et al. 2020).

## ACKNOWLEDGEMENTS

We thank Professor Christopher K. Glass for the gift of PWK/PhJ breeders. We thank Wilfredo Villanueva and Abraham Ramirez for excellent animal care. W.A.A. was supported in part by an institutional training grant to the UC San Diego Genetics Training Program (T32 GM008666) from the National Institute of General Medical Sciences and by a Ruth L. Kirschstein National Research Service Award (F31 NS061513) from the National Institute of Neurological Disorders and Stroke. This work was supported by grants R01 NS054871 and R01 NS097534 from the National Institute of Neurological Disorders and Stroke to B. A. H.

## SUPPLEMENTAL FILES

**Supplemental Data S1**: B6 x BALB/c intercross data for Figure 3A, C. Data are in “csvr” format for R/qtl.

**Supplemental Data S2**: B6 x BALB/c backcross data for Figure 3B, D. Data are in “csvr” format for R/qtl.

**Supplemental Table S1**: Fisher’s exact test p-values for pairwise comparisons of *Zfp423*^*nur12*^ survival frequency data among B6 backcrosses from 13 strains, with false-discovery rate corrections for 78 comparisons.

**Supplemental Table S1.** Pairwise tests for difference in survival frequency.

| Strain1 | Strain2 | FisherExact-p | FDR-adjusted-p |
| --- | --- | --- | --- |
| PWK | 129S1 | 0.7764 | 0.8296 |
| PWK | FVB | 0.1354 | 0.2515 |
| PWK | BALB | 0.007827 | 0.0226 |
| PWK | NOD | 0.003394 | 0.0110 |
| PWK | C3H | 0.0007257 | 0.0029 |
| PWK | DBA | 0.02394 | 0.0584 |
| PWK | NZO | 0.0007466 | 0.0029 |
| PWK | C57L | 1.12E-06 | 7.94E-06 |
| PWK | C57BLKS | 4.24E-08 | 6.61E-07 |
| PWK | C58 | 4.71E-09 | 1.14E-07 |
| PWK | AJ | 3.12E-09 | 1.14E-07 |
| PWK | BTBR | 3.14E-07 | 3.32E-06 |
| 129S1 | FVB | 0.2022 | 0.3584 |
| 129S1 | BALB | 0.01639 | 0.0412 |
| 129S1 | NOD | 0.005928 | 0.0178 |
| 129S1 | C3H | 0.001363 | 0.0048 |
| 129S1 | DBA | 0.04187 | 0.0961 |
| 129S1 | NZO | 0.001215 | 0.0045 |
| 129S1 | C57L | 1.72E-06 | 1.12E-05 |
| 129S1 | C57BLKS | 5.68E-08 | 7.38E-07 |
| 129S1 | C58 | 5.87E-09 | 1.14E-07 |
| 129S1 | AJ | 3.84E-09 | 1.14E-07 |
| 129S1 | BTBR | 6.63E-07 | 5.17E-06 |
| FVB | BALB | 0.7696 | 0.8296 |
| FVB | NOD | 0.3365 | 0.5102 |
| FVB | C3H | 0.0892 | 0.1739 |
| FVB | DBA | 0.2797 | 0.4452 |
| FVB | NZO | 0.05442 | 0.1147 |
| FVB | C57L | 0.004633 | 0.0145 |
| FVB | C57BLKS | 0.001909 | 0.0065 |
| FVB | C58 | 0.0005045 | 0.0023 |
| FVB | AJ | 0.0003496 | 0.0017 |
| FVB | BTBR | 0.0006019 | 0.0026 |
| BALB | NOD | 0.2531 | 0.4200 |
| BALB | C3H | 0.04588 | 0.1022 |
| BALB | DBA | 0.3467 | 0.5102 |
| BALB | NZO | 0.04718 | 0.1022 |
| BALB | C57L | 0.0001376 | 7.16E-04 |
| BALB | C57BLKS | 6.60E-06 | 3.96E-05 |
| BALB | C58 | 4.08E-07 | 3.54E-06 |
| BALB | AJ | 3.40E-07 | 3.32E-06 |
| BALB | BTBR | 6.59E-05 | 3.67E-04 |
| NOD | C3H | 0.5264 | 0.6562 |
| NOD | DBA | 0.7686 | 0.8296 |
| NOD | NZO | 0.3163 | 0.4934 |
| NOD | C57L | 0.06793 | 0.1394 |
| NOD | C57BLKS | 0.02907 | 0.0687 |
| NOD | C58 | 0.01301 | 0.0350 |
| NOD | AJ | 0.009808 | 0.0273 |
| NOD | BTBR | 0.01453 | 0.0378 |
| C3H | DBA | 1 | 1.0000 |
| C3H | NZO | 0.7694 | 0.8296 |
| C3H | C57L | 0.4711 | 0.6228 |
| C3H | C57BLKS | 0.3434 | 0.5102 |
| C3H | C58 | 0.2184 | 0.3734 |
| C3H | AJ | 0.1239 | 0.2357 |
| C3H | BTBR | 0.08217 | 0.1643 |
| DBA | NZO | 0.6968 | 0.8112 |
| DBA | C57L | 0.4671 | 0.6228 |
| DBA | C57BLKS | 0.4421 | 0.6050 |
| DBA | C58 | 0.4062 | 0.5867 |
| DBA | AJ | 0.2202 | 0.3734 |
| DBA | BTBR | 0.1606 | 0.2913 |
| NZO | C57L | 1 | 1.0000 |
| NZO | C57BLKS | 0.7653 | 0.8296 |
| NZO | C58 | 0.5384 | 0.6562 |
| NZO | AJ | 0.5027 | 0.6428 |
| NZO | BTBR | 0.4277 | 0.5957 |
| C57L | C57BLKS | 1 | 1.0000 |
| C57L | C58 | 0.6717 | 0.7938 |
| C57L | AJ | 0.5017 | 0.6428 |
| C57L | BTBR | 0.2732 | 0.4440 |
| C57BLKS | C58 | 0.8442 | 0.8780 |
| C57BLKS | AJ | 0.537 | 0.6562 |
| C57BLKS | BTBR | 0.4155 | 0.5893 |
| C58 | AJ | 0.8329 | 0.8779 |
| C58 | BTBR | 0.5728 | 0.6874 |
| AJ | BTBR | 0.7618 | 0.8296 |

## Notes

### Competing Interest Statement

The authors have declared no competing interest.

### Summary of Updates

Added co-author Alison Ponn. Revised text for clarity, especially in materials and methods. Added supplemental table of nominal and FDR-corrected p-values for comparisons of survival frequencies among backcrosses to different mapping strains.

